# Pan-Coronavirus Protease Engagement and Differentiated Pharmacology of Antiviral Candidate CMX990

**DOI:** 10.64898/2026.09.29.755428

**Authors:** N. G. R. Dayan Elshan, Karen C. Wolff, Laura Riva, Malina A. Bakowski, Simeon Babarinde, Anshul Chaudhary, Mark Zeller, Ashley K. Woods, Gennadii Grabovyi, Katy B. Wilson, Rebecca A. Leonard, Briana L. McGovern, Jared Benjamin, Isidora Suazo, Mary Gordon, Frank O. Weiss, Sourav Ghorai, James R. Pedroarena, Kristi L. Marquardt, Maheeka Bimal, Stuart Weston, Carly Dillen, Appaso M. Jadhav, Melanie G. Kirkpatrick, Amal Gebara-Lamb, Nisar Farhat, Vân T. B. Nguyen-Tran, Victor Chi, Kyoung-Jin Lee, Case W. McNamara, Anil Kumar Gupta, Alireza Rahimi, John R. Teijaro, Jian Jeffrey Chen, Sean B. Joseph, Matthew Frieman, Nicholas S. Heaton, Kris White, Ian A. Wilson, Peter G. Schultz, Sumit K. Chanda, Arnab K. Chatterjee

## Abstract

COVID-19 highlighted the need for antivirals that can be rapidly deployed against both current and future coronavirus threats. CMX990 is a clinical stage coronavirus main protease (M^pro^) inhibitor bearing a unique trifluoromethoxymethyl ketone warhead. Here, we broadly evaluate its pan-coronavirus target engagement, antiviral breadth, and pharmacologic differentiation. Across a reporter panel of 24 M^pro^ enzymes spanning alpha-, beta-, delta-, and gamma-coronaviruses, CMX990 retained robust potent activity, including against nirmatrelvir-resistant SARS-CoV-2 variants. In direct comparison, nirmatrelvir and ensitrelvir showed diminished potency against non-SARS-CoV-2 coronaviruses. Live-virus assays corroborated this breadth, with potent, selective activity against SARS-CoV-2, SARS-CoV-1, MERS-CoV, and additional human coronaviruses. Structural studies revealed a conserved covalent binding mode centered on the catalytic Cys of coronavirus M^pro^. In serial-passage studies, resistance emerged only after approximately twice as many serial passages under CMX990 selection as under nirmatrelvir selection, without selecting for the canonical nirmatrelvir-resistance substitution E166V. Extracellular washout experiments revealed superior persistence of antiviral activity relative to nirmatrelvir, consistent with durable target engagement. In a murine SARS-CoV-2 infection model, once-daily CMX990 protected against weight loss and reduced lung viral titers to near baseline levels. Collectively, these findings identify CMX990 as a structurally and functionally validated pan-coronavirus protease inhibitor supporting its potential as a pandemic-ready antiviral candidate.

## Introduction

Coronaviruses (CoVs) have repeatedly demonstrated their capacity to cause large-scale human outbreaks, including the severe acute respiratory syndrome (SARS) epidemic in 2002–2003 (SARS-CoV-1), Middle East respiratory syndrome (MERS) beginning in 2012, and the coronavirus disease 2019 (COVID-19) pandemic caused by SARS-CoV-2.^1^ Together, these events highlight the persistent risk posed by zoonotic coronaviruses and underscore the need for broadly active antiviral therapeutics that can be rapidly deployed against both current and future emergent strains. The COVID-19 pandemic alone resulted in hundreds of millions of infections worldwide and emphasized the vulnerability of global health systems to novel viral pathogens. While vaccines have played a central role in controlling disease burden, their effectiveness can be diminished by viral evolution, manufacturing constraints, and unequal global distribution, highlighting the continued importance of small-molecule antivirals as a complementary and resilient countermeasure.^2, 3^

Among the viral targets explored for coronavirus therapeutics, the main protease (M^pro^, also referred to as 3CL^pro^) has emerged as one of the most attractive and clinically validated drug targets. M^pro^ mediates the proteolytic processing of viral polyproteins required for replication and transcription and is indispensable for the viral life cycle. Importantly, the active site architecture and substrate-binding pockets of M^pro^ are highly conserved across diverse coronaviruses, including SARS-CoV-2, SARS-CoV-1, and MERS-CoV, enabling the possibility of developing inhibitors with activity spanning multiple coronavirus lineages. The clinical success of M^pro^ inhibitors, such as nirmatrelvir, has validated this target class. However, currently available therapeutics were primarily optimized for SARS-CoV-2 and do not necessarily address the broader challenge of future coronavirus emergence.

The development of pan-coronavirus antivirals represents an increasingly important strategy for pandemic preparedness. Because zoonotic coronaviruses continue to circulate in animal reservoirs, future spillover events remain likely.^4^ Broad-spectrum antivirals targeting conserved viral machinery could provide immediate therapeutic options during the early phases of emerging outbreaks, before pathogen-specific countermeasures can be developed. Recent efforts have therefore focused on identifying inhibitors capable of engaging the conserved catalytic machinery of M^pro^ across multiple coronavirus clades. Such agents could function both as frontline treatments for current infections and as part of a pre-positioned antiviral arsenal against future coronavirus threats. We previously reported the discovery of CMX990 (**Figure 1**), a potent small-molecule inhibitor of SARS-CoV-2 M^pro^ bearing a trifluoromethoxymethyl ketone warhead.^5^ CMX990 demonstrated potent inhibition of SARS-CoV-2 replication and differentiated pharmacokinetic properties relative to existing agents, including improved *in vitro* metabolic stability and the potential for unboosted oral dosing.

**Figure 1.**
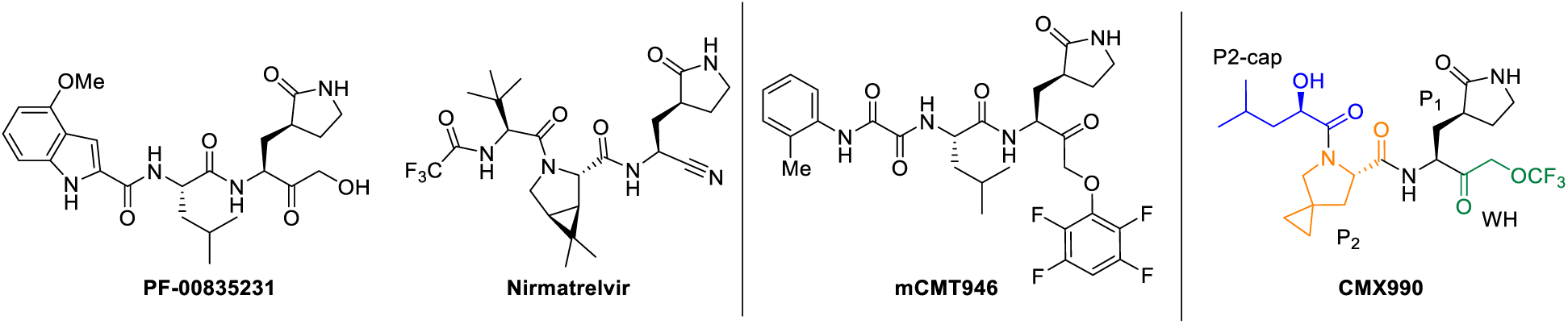
CMX990 and the comparator compounds utilized in this work. CMX990 is a clinical-stage coronavirus main protease inhibitor with unique broad-spectrum pharmacological efficacy. Structurally, it features a trifluoromethoxymethyl ketone warhead (WH), a P1 amino acid motif bearing an enzyme recognition motif, a P2 proline substituent, and a leucic acid P2-*N*-cap. Described compounds represent varying degrees of warhead reactivities. The aryloxy group bearing mCMT946 is anticipated to be the most reactive, while the hydroxymethylketone (e.g., PF-00835231) and nitrile (nirmatrelvir) warheads are anticipated to have the lowest reactivity.^6^

In the current work, we comprehensively define the pan-coronavirus protease engagement and the differentiated pharmacology of CMX990, disclosing data from multiple complementary *in vitro* and *in vivo* assessment methods, including protease reporter panels, structural biology studies, expanded cell-based antiviral assays, resistance-selection, washout studies, and a proof-of-concept efficacy study. Together, these studies establish CMX990 as a structurally and functionally validated pan-coronavirus protease inhibitor and highlight the broader feasibility of targeting conserved coronavirus protease architecture to enable pandemic-ready antiviral therapeutics.

## Results and Discussion

### CMX990 has potent broad-spectrum reporter, biochemical, and cell-based activity against coronaviruses

To evaluate the breadth of coronavirus main protease inhibition in a cellular context, CMX990 was profiled alongside nirmatrelvir and ensitrelvir in a reporter panel^7^ representing 24 M^pro^ enzymes from alpha-, beta-, delta-, and gamma-coronaviruses, including human pathogens, animal viruses with zoonotic potential, and SARS-CoV-2 variants associated with reduced susceptibility to nirmatrelvir (**Figure 2**). Reporter activity was measured after treatment with each inhibitor at 10 μM and normalized to the uninhibited condition; thus, lower residual reporter signal indicates more complete protease inhibition. CMX990 produced near-complete inhibition of wild-type SARS-CoV-2 M^pro^, leaving 1.3% residual signal compared with 12.7% and 38.3% for nirmatrelvir and ensitrelvir, respectively. Importantly, this activity was maintained against the SARS-CoV-2 M^pro^ variants S144A, P132H, and L50F/E166V (nirmatrelvir-resistant mutant^8^), for which CMX990 yielded only 0.6 – 1.7% residual signal. In contrast, the corresponding residual signals for nirmatrelvir ranged from 32.7% to 100%, with the largest loss of activity observed against L50F/E166V, whereas ensitrelvir left 50.5 – 80.8 % residual activity. These findings indicate that CMX990 engages both wild-type and inhibitor-resistant SARS-CoV-2 proteases more effectively than the tested comparator agents under identical assay conditions.

**Figure 2.**
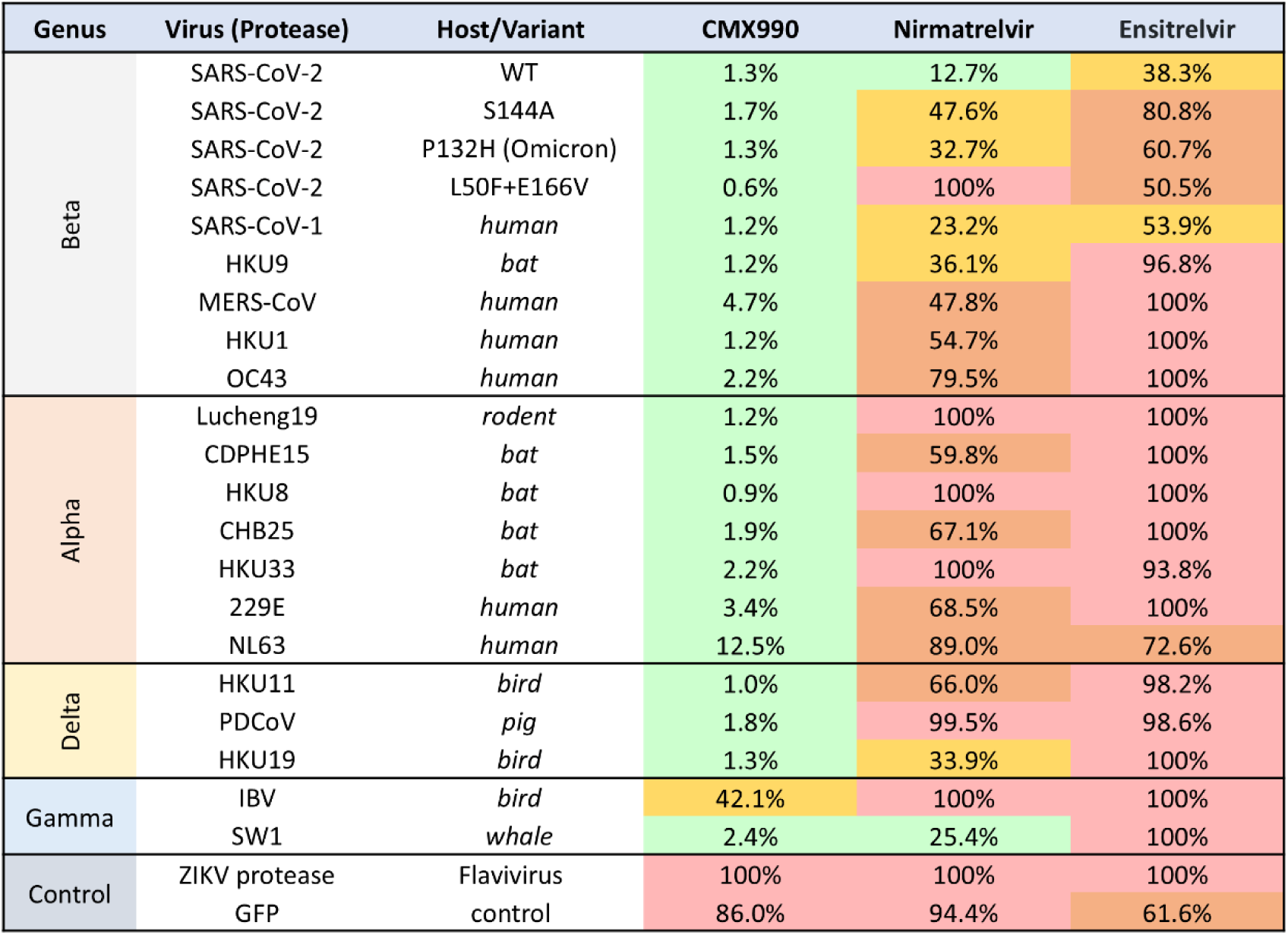
M^pro^ inhibitor CMX990 has superior pan-coronavirus inhibition potential as compared to nirmatrelvir and ensitrelvir at identical conditions. Normalized cell-based M^pro^ reporter signal is shown as % of non-inhibited signal (DMSO), after treatment with each protease inhibitor (10 µM). Better inhibition results in lower reporter signal (green color coded). CMX990 shows superior pan-activity against all tested coronavirus proteases (including from other species) and drug-resistant mutants (e.g., L50F+E166V). Representative data were also confirmed in live-virus assays.

The breadth of CMX990 extended well beyond SARS-CoV-2. Among other betacoronaviruses, including SARS-CoV-1, MERS-CoV, HKU1, OC43, and a bat HKU9 protease, CMX990 restricted residual reporter activity to 1.2 – 4.7%. By comparison, nirmatrelvir left 23.3 – 79.5% residual signal and ensitrelvir 53.9 – 100%, demonstrating progressively reduced engagement of proteases more distantly related to SARS-CoV-2. A similarly differentiated profile was observed across the alphacoronavirus panel, which included rodent, bat, and human viruses: CMX990 generally reduced reporter activity to less than 4%, with the exception of NL63 M^pro^, for which 12.5% residual signal remained. Nirmatrelvir and ensitrelvir were substantially less active across this group, frequently leaving more than 60% residual activity and, for several proteases, producing values near or above uninhibited control. CMX990 also strongly inhibited each delta coronavirus protease examined, leaving only 1.0 – 1.8% residual signal, whereas the comparator inhibitors showed markedly weaker to no engagement. Within the gamma coronavirus group, CMX990 inhibited the whale coronavirus SW1 protease to 2.4% residual activity but was less active against infectious bronchitis virus M^pro^, for which 42.1% reporter activity remained, identifying a potential boundary to its otherwise broad coverage. The absence of appreciable inhibition of the unrelated Zika virus protease supports the coronavirus-protease selectivity of the observed profile. Collectively, the reporter panel demonstrates that CMX990 maintains unusually broad intracellular engagement across phylogenetically diverse coronavirus M^pro^ enzymes and retains activity against protease variants that compromise nirmatrelvir and ensitrelvir. This profile provides a mechanistic basis for the broad antiviral activity observed in subsequent live-virus assays and supports the potential value of CMX990 as a pre-positioned antiviral for future coronavirus emergence.

Viral replication assays (**Table 1**) corroborated the above broad-engagement findings (**Figure 2**), where CMX990 potently, but selectively inhibited viral growth (CC_50_ > 40 µM). Importantly, it was not as active against viruses that lack homologous targets to SARS-CoV-2 M^pro^. As described previously^5^, CMX990 is generally equal or better in potency (>4× viral yield reduction, VYR) to nirmatrelvir, in biochemical potency versus SARS-CoV-2 M^pro^, and in live virus assays of wild type and variant forms of SARS-CoV-2. Here, we extend the profiling data to additional coronaviruses and to additional viral families, to further establish the pan-antiviral breadth as well as selectivity of CMX990. Potency values in **Table 1** were generated across multiple cell types, readouts and assay endpoints (cytopathic effect, CPE; immunofluorescence, IF; viral yield reduction, VYR). Where multiple measurement types are available (e.g., SARS-CoV-2, SARS-CoV-1, and MERS-CoV), the derived values are directionally consistent

**Table 1.**
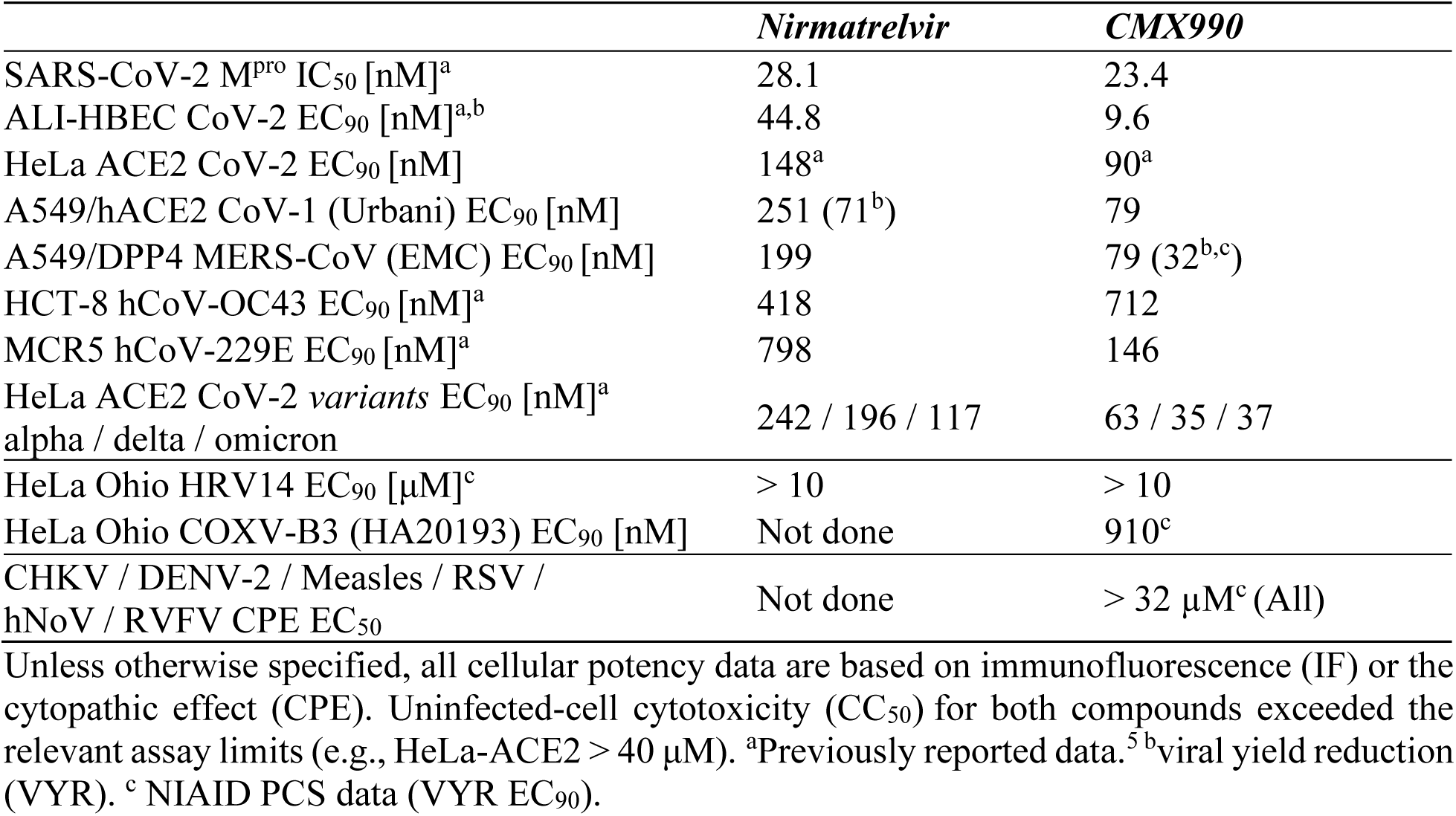
***In vitro* antiviral potency profile of CMX990.**

Against SARS-CoV-1 (Urbani strain) in A549-hACE2 cells, CMX990 was approximately 3-fold more potent than nirmatrelvir (EC_90_ 79 nM vs. 251 nM). Against MERS-CoV (EMC strain) in A549/DPP4 cells, CMX990 again outperformed nirmatrelvir by a comparable margin (79 nM vs. 199 nM). To further assess selectivity, CMX990 was tested against viruses from other viral families (**Figure 2**). When tested versus enteroviruses, CMX990 showed modest potency against coxsackievirus B3 (EC_90_ 910 nM); but both CMX990 and nirmatrelvir showed no appreciable activity against human rhinovirus 14 (HRV14, EC_90_ > 10 µM). The modest activity versus coxsackievirus B3 is attributed to the chymotrypsin-like cysteine-protease fold derived shared homology between the picornavirus 3C protease and coronavirus M^pro^. Against a broader panel of phylogenetically unrelated RNA viruses lacking an M^pro^ ortholog (chikungunya, dengue-2, measles, RSV, human norovirus, and Rift Valley fever virus), CMX990 showed no significant antiviral activity at concentrations up to 32 µM, supporting mechanism-specific antiviral activity of CMX990 rather than general antiviral or cytotoxic activity.

To evaluate the selectivity of CMX990 against human proteases, the compound was profiled in a broad mammalian peptidase panel (Eurofins; **Table 2**). CMX990 exhibited minimal activity against the majority of proteases evaluated (IC_50_ > 10 μM) despite its peptidomimetic scaffold. The principal off-target activities were limited to cathepsins K, L, L2 (V), and S, while no substantial inhibition of other cysteine, serine, aspartyl, or metalloproteases was observed. Some degree of cathepsin inhibition is expected for peptide-derived coronavirus M^pro^ inhibitors because both enzyme families are cysteine proteases that recognize extended peptide substrates and share related active-site architectures. Indeed, nirmatrelvir also inhibits cathepsins, particularly cathepsins K and S, at pharmacologically relevant concentrations (**Table 2**). Given the acute treatment windows for coronavirus protease inhibitor use, the anticipated risks from these off-target engagement of select cathepsins is limited. Importantly, this profile is not attributed to the trifluoromethoxymethyl ketone warhead itself, where structurally related analogs (not shown) bearing the same electrophilic warhead display up to approximately 50-fold differences in cathepsin inhibition, indicating that off-target cathepsin engagement is primarily dictated by the overall scaffold rather than intrinsic warhead reactivity.

**Table 2.**
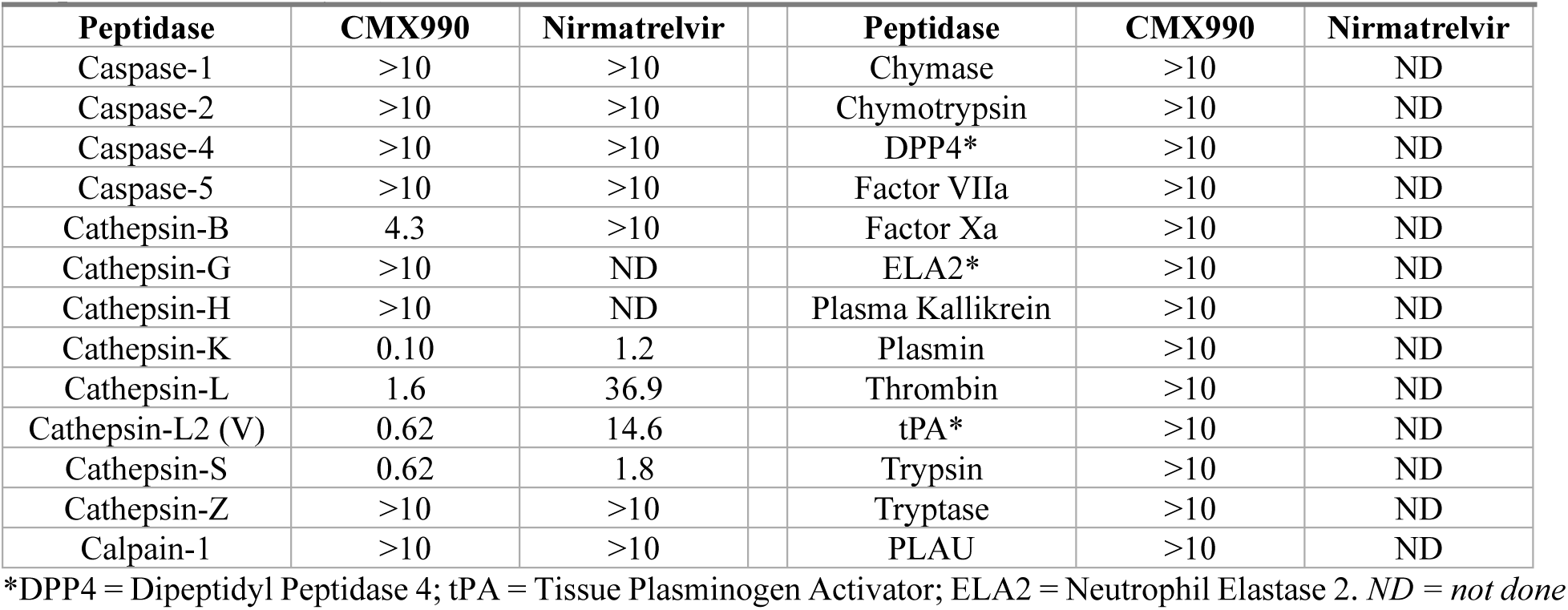
Mammalian peptidase inhibition data for CMX990 vs nirmatrelvir. (Potencies expressed as IC_50_; µM)

### Target engagement of CMX990 further corroborated via structural biology

To structurally rationalize the broad pan-coronavirus engagement, we determined co-crystal structures of CMX990 bound to SARS-CoV-1, SARS-CoV-2, and MERS-CoV M^pro^ (**Figure 3**). In the SARS-CoV M^pro^ homodimer, CMX990 occupies the substrate-binding cleft of each protomer (**Figure 3a**), consistent with engagement of the enzyme’s catalytically competent dimeric form. A close-up view of the binding site (**Figure 3b**) shows that the inhibitor forms a covalent bond to the catalytic nucleophile Cys145, with the electrophilic warhead positioned directly adjacent to His41, the second member of the Cys-His catalytic dyad. Beyond this covalent linkage, CMX990 is stabilized by an extended hydrogen-bonding network involving the side chain of His163 and backbone atoms of Phe140, Gly143, Cys145, His164, and Glu166, together with an additional interaction involving the Glu166 side chain. Notably, several of these interactions recapitulate conserved contacts involved in recognition of the P1 glutamine within the S1 substrate-binding pocket of M^pro^.^9^

**Figure 3.**
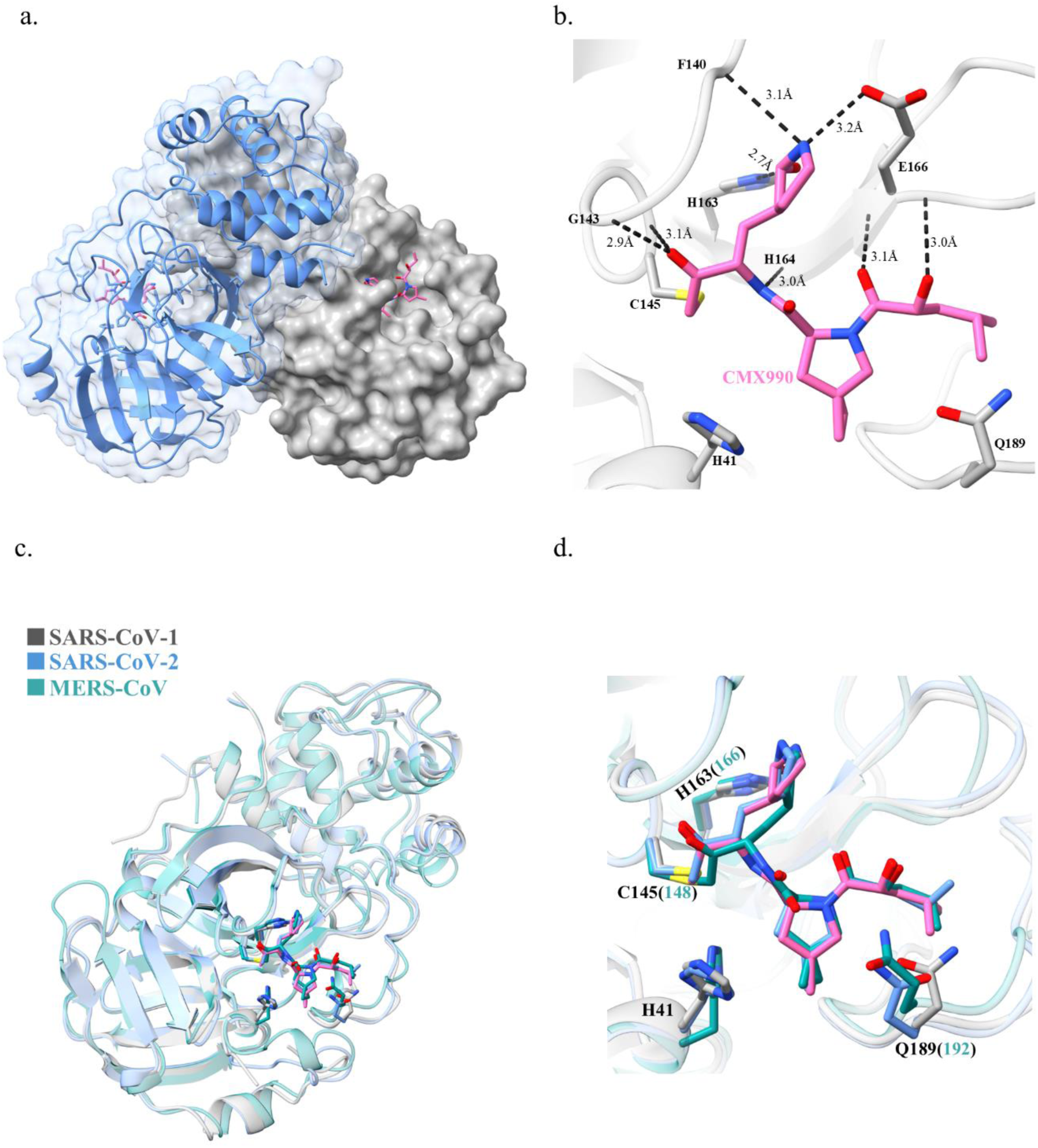
Structural basis for broad-spectrum inhibition of coronavirus main proteases by CMX990. (a) SARS-CoV-1 M^pro^ homodimer (gray surface / blue cartoon) in complex with CMX990 (pink sticks) that occupies the substrate-binding cleft of each protomer. (b) Close-up view of the SARS-CoV-1 M^pro^-CMX990-binding site. The inhibitor forms a covalent bond with catalytic Cys145 and is stabilized by hydrogen-bonding interactions (black dashed lines) with the side chain of His163, the backbone atoms of Phe140, Gly143, Cys145, His164, and Glu166, together with an additional interaction to Glu166 side chain. The catalytic His41 is also shown adjacent to CMX990. (c) Structural superposition of the SARS-CoV-1 (gray), SARS-CoV-2 (blue), and MERS-CoV (turquoise) M^pro^–CMX990 complexes. The three proteases adopt highly similar overall structures despite some sequence divergence, with the inhibitor occupying a conserved position within the substrate-binding cleft. (d) Superposition of the active sites of SARS-CoV-1, SARS-CoV-2, and MERS-CoV M^pro^ complexes. The catalytic residues and surrounding substrate-binding pocket adopt highly conserved conformations, resulting in a nearly identical binding orientation of CMX990 across the three coronavirus proteases despite differences in their active-site sequences (**Figure S1**).

Superposition of the SARS-CoV-1, SARS-CoV-2, and MERS-CoV complexes (**Figure 3c-d**) shows that despite sequence divergence between these three virus lineages, the overall M^pro^ fold and, in particular, the local geometry of the catalytic dyad and surrounding substrate-binding pocket is highly conserved. As a result, CMX990 adopts a nearly identical binding pose and covalent attachment geometry across all three proteases (**Figure 3d**). This structural conservation provides a mechanistic basis for the broad inhibition observed across the coronavirus panel in **Figure 2** and directly rationalizes the retained potency across SARS-CoV-2, SARS-CoV-1, and MERS-CoV in **Table 1**. Rather than relying on lineage-specific contacts, CMX990 engages residues that are under strong evolutionary constraint because they are required for the catalytic function of M^pro^, as well as multiple interactions to the M^pro^ backbone.

### CMX990 exhibits sustained antiviral activity following compound washout

The aforementioned structural data also inform upon the compound’s kinetic behavior. To determine whether CMX990’s covalent engagement translates into a durable antiviral effect after compound clearance, and to help delineate mechanistic differences among inhibitor classes, we performed a compound washout study in a HeLa-ACE2 single-cycle infection assay (**Figure 4**). The covalent bond to Cys145 is consistent with the sustained antiviral effect after compound washout (**Figure 4**), in which CMX990 continued to suppress viral RNA production well after extracellular compound had been removed, tracking between the irreversible internal control mCMT946 (**Figure 1**) and the reversibly binding nirmatrelvir. Cells were synchronously infected 1 hr at 4°C in the presence of the compound of interest [10× EC_50_]. Inoculum was then removed and fresh compounds were added, followed by incubation at 37 °C. Media with compounds were then removed at 2, 4, or 6 h post-infection, replaced by assay media, and viral RNA was quantified at 10 h; a no-washout condition (compound present throughout the experiment) served as the maximal-inhibition reference. The panel included the covalent protease inhibitors CMX990 and mCMT946 (an internal irreversible-inhibitor control, with a tetrafluorophenoxy warhead), the reversible oral benchmark nirmatrelvir, and two mechanistically distinct comparators – the nucleoside analog remdesivir and the PIKfyve/entry inhibitor apilimod - to help contextualize the protease-inhibitor washout kinetics.

**Figure 4.**
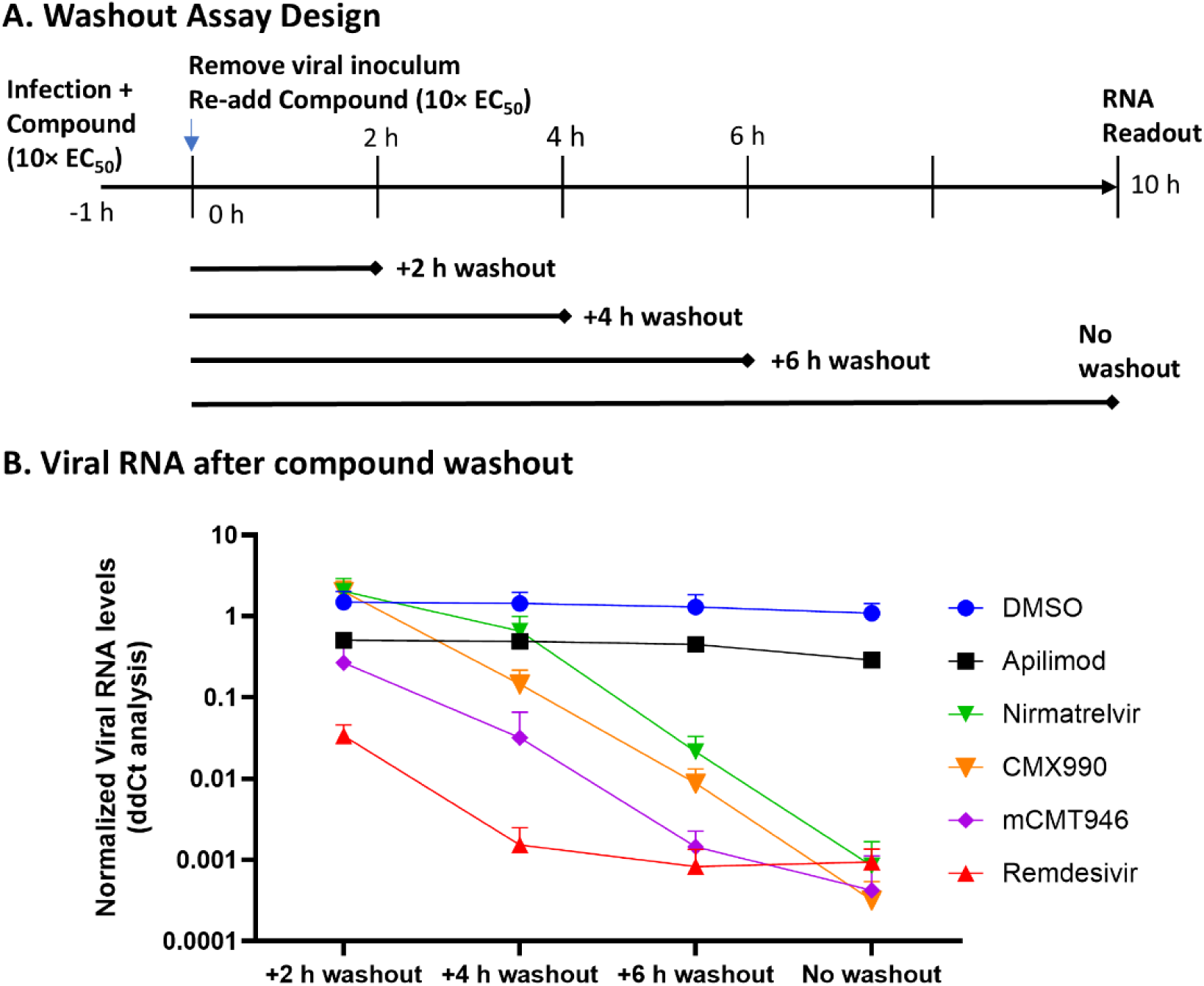
Compound washout experiments reveal greater persistence of antiviral activity for CMX990 than nirmatrelvir. (A) Washout assay design: SARS-CoV-2–infected HeLa-ACE2 cells were dosed with each compound at 10× its own EC_50_ at the time of infection (T = -1 h); viral inoculum was removed after one hour by washing the cell monolayers (100 µL × 2 ice-cold PBS), and compounds diluted in assay media were freshly added to each well (T = 0 h); compound was removed by washout at 2, 4, or 6 h, or retained throughout (no washout), and viral RNA was quantified at 10 h. (B) Normalized viral RNA levels across washout conditions for DMSO, apilimod, remdesivir, nirmatrelvir, mCMT946 (internal irreversible control with a tetrafluorophenoxymethyl ketone warhead), and CMX990. CMX990 shows a greater reduction in viral RNA than nirmatrelvir at the 4 h washout time point, consistent with durable covalent M^pro^ engagement. Apilimod (an entry-stage inhibitor) and remdesivir (a nucleoside analog requiring intracellular triphosphate conversion) are included as mechanistic reference points and are discussed further in the text. Each data point represented as an arithmetic mean of three individual experiments with three technical replicates within each plate.

Apilimod behaved as expected for an entry-stage inhibitor, maintaining strong suppression of viral RNA essentially independent of washout time. Because apilimod blocks a step early in the viral life cycle rather than a continuously required enzymatic activity, its effect does not depend on sustained intracellular compound residence in the same way a protease inhibitor does, and therefore serves as a useful mechanistic reference point for interpreting the protease inhibitor curves. Remdesivir was more potent in this assay than anticipated, showing near-maximal suppression even at the earliest (2 h) washout point. We suspect this reflects the pharmacology of the nucleoside class rather than unusually slow target dissociation: remdesivir must be converted intracellularly to its active triphosphate metabolite, and that metabolite pool likely persists after the parent compound is washed out of the extracellular medium, confounding this assay as a direct readout of durable target engagement for this compound class.

Among the protease inhibitors, clear differentiation was observed. mCMT946, the irreversible internal control, showed the greatest suppression of viral RNA upon washout at every time point tested, consistent with its highly reactive irreversibly covalent warhead – leading to non-dissociable target engagement. CMX990 showed a somewhat less pronounced, but still substantial, washout effect than mCMT946, and outperformed nirmatrelvir at each washout time point, most notably at 4 h, where CMX990 suppressed viral RNA by roughly an order of magnitude more than nirmatrelvir under identical removal conditions. By 6 h washout, CMX990 approached the suppression seen in the no-washout condition, indicating that once bound, it continues to restrict viral replication well after free compound is no longer present – behavior consistent with durable, covalent occupancy of the M^pro^ active site (as supported independently by the structural data in **Figure 3**), and in contrast to the reversibly-covalent nirmatrelvir.

From a translational standpoint, this persistence of antiviral activity after washout could be particularly relevant in tissue compartments where exposure transiently dips below the EC_90_ threshold between doses, a scenario in which a compound with fast off-kinetics would lose antiviral coverage while a durably-engaged inhibitor like CMX990 would be expected to retain effect. The magnitude of the washout effect also appears to have SAR traction across the warhead and P2/P2-cap positions, suggesting durability of target engagement, and by extension pharmacodynamic advantage, could potentially be further optimized in follow-up analogs.

### CMX990 shows a superior resistance profile versus comparators in live-virus assays

CMX990 exhibited a differentiated resistance profile compared to nirmatrelvir and PF-00835231 in live-virus serial-passage studies (**Figure 5**). The delayed resistance phenotype may reflect the combination of durable covalent engagement and the conserved interaction network formed by CMX990. However, the relative contribution of covalent kinetics versus scaffold-specific interactions cannot be exclusively resolved from these experiments. SARS-CoV-2 (WA1/2020) was passaged in HeLa-ACE2 cells under increasing concentrations of inhibitor treatment, and resistance emergence was tracked as the passage number at which virus actively replicated at 32× the parental EC_50_. Resistance to nirmatrelvir emerged by passage 8, and to PF-00835231 by passage 9, whereas virus resistant to CMX990 did not emerge until passage 16 - roughly double the number of passages required for the comparator protease inhibitors (**Figure 5A**).

**Figure 5.**
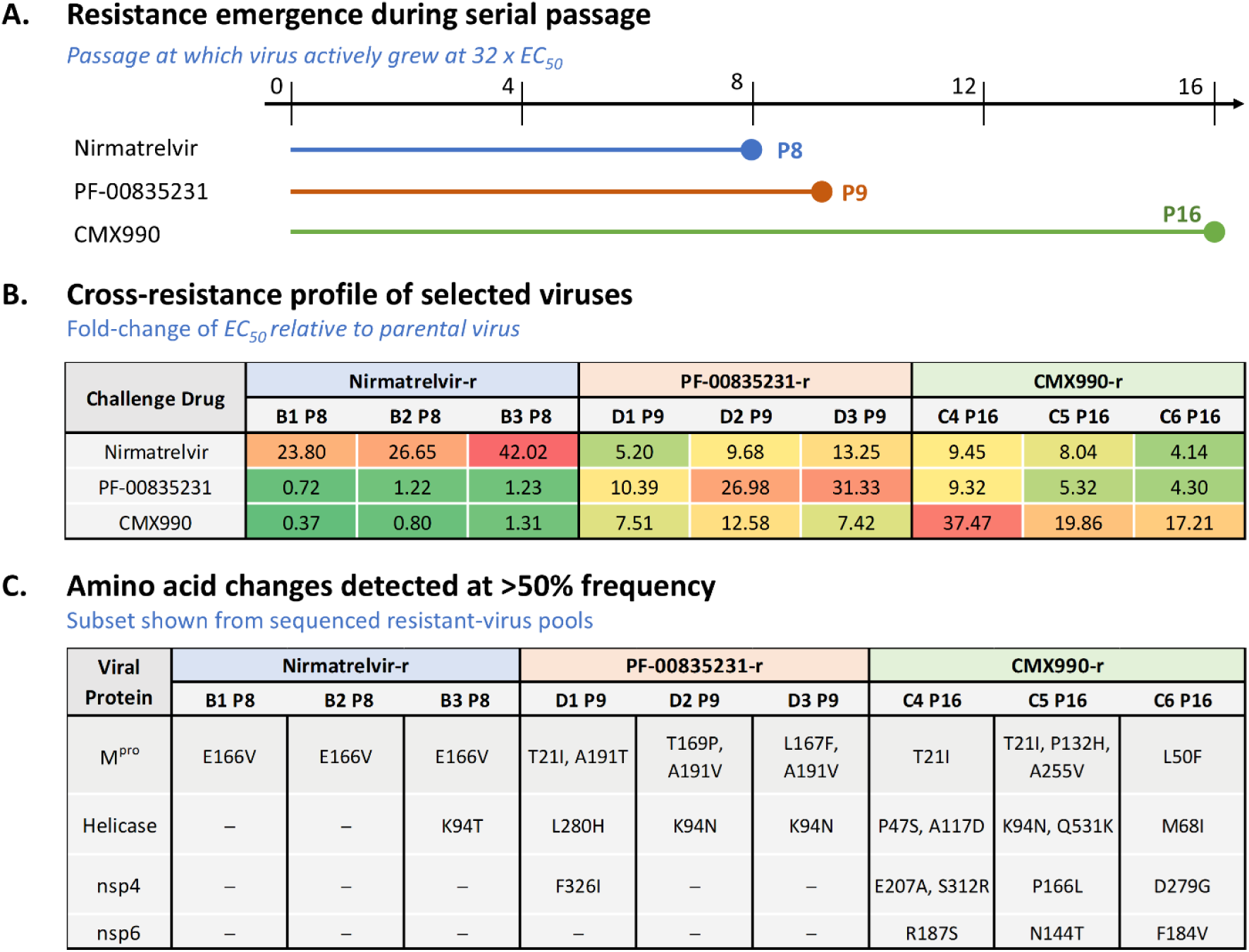
Resistance-selection yielded a differentiated profile for CMX990 relative to nirmatrelvir and PF-00835231. (A) Passage number at which SARS-CoV-2 (WA1/2020) actively replicated under selective pressure from each inhibitor (32× the parental EC₅₀) during serial passage in HeLa-ACE2 cells. Resistant virus emerged at passage 8 for nirmatrelvir, passage 9 for PF-00835231, and passage 16 for CMX990. (B) Cross-resistance profile of viral populations selected under nirmatrelvir (wells B1–B3, passage 8), PF-00835231 (wells D1–D3, passage 9), and CMX990 (wells C4–C6, passage 16) pressure, shown as EC₅₀ fold-change relative to parental virus upon challenge with each of the three inhibitors. (C) Amino acid substitutions detected at >50% frequency by sequencing of the resistant viral pools shown in (B), reported for Mpro and selected nonstructural proteins (helicase, nsp4, nsp6).

Cross-resistance testing of the selected viral populations revealed a clear asymmetry among the three inhibitors (**Figure 5B-C**). Virus selected for resistance to nirmatrelvir converged on the previously reported M^pro^ E166V substitution^10–13^ in all three replicates and was not cross-resistant to either CMX990 nor PF-00835231, indicating that E166V strongly compromises nirmatrelvir while retaining susceptibility to CMX990 and PF-00835231 under these assay conditions. Virus selected for resistance to PF-00835231 acquired multiple combinations of M^pro^ substitutions (T21I/A191T, T169P/A191V, L167F/A191V) that showed some cross-resistance to both nirmatrelvir and CMX990 (**Figure 5B**), indicating a broader impact resistance mechanism than the nirmatrelvir driven mutation of E166V.

Virus selected under CMX990 pressure also showed cross-resistance to both comparator compounds. Here, a comparatively large number of amino acid changes were detected across the viral genome (M^pro^, helicase, nsp4, and nsp6), and these did not correlate cleanly with the degree of phenotypic resistance. Several of the observed substitutions (T21I, L50F, P132H) are known to occur commonly in circulating or passaged virus independent of drug pressure,^14^ and CMX990-resistant virus notably did not select for the E166V substitution that drives nirmatrelvir resistance. The changes detected in nsp4, nsp6, helicase, and (in some replicates) the surface glycoprotein may reflect genuine contributors to reduced susceptibility or may simply represent compensatory adaptations to serial passage under drug pressure in vitro rather than mutations that directly reduce CMX990 potency. These data are consistent with the findings of Lewandowski et al.,^14^ where mutations distal to the active site were shown important to confer viral resistance. In that work, the non-active site mutations L50F, T21I, and P252L restored the fitness of viral replication. This effect was mapped to the impact of these residues in enhancing the M^pro^ dimer-dimer interface interactions and to helping place the nsp5-6 substrate at the enzyme catalytic center.^14^ This observation is consistent with the retained potency of CMX990 against the L50F/E166V and P132H variants in the reporter panel **(Figure 2**). Mapping these M^pro^ variants onto the SARS-CoV-2 M^pro^-CMX990 structure showed that T21, L50, P132, and T169 are distal to the bound inhibitor, whereas A191 resides in the second shell and L167 is positioned at the edge of the binding pocket. In contrast, E166 directly contacts CMX990. Notably, CMX990 also makes contacts with the E166 backbone in both protomers, which would be retained upon substitution of E166 side chain, and may contribute to the retained activity.

### CMX990 shows proof-of-concept efficacy in a SARS-CoV-2 mouse model

To determine whether the pan-coronavirus target engagement, resistance barrier, and target-residence advantages described above translate into in vivo antiviral benefit, we evaluated CMX990 in a murine SARS-CoV-2 infection model (**Figure 6**). Female 129S1 mice were intranasally infected with mouse-adapted SARS-CoV-2 and treated beginning 1 h before infection with either vehicle, nirmatrelvir (600 mg/kg, PO, BID), or CMX990 (100 mg/kg, SC, QD), with dosing continued once daily for 3 days (**Figure 6A**). CMX990 was dosed subcutaneously rather than orally for this study because, as we reported previously,^5^ CMX990 has lower oral exposure in mice than nirmatrelvir; the subcutaneous route was therefore selected to ensure adequate systemic exposure (**Figure 6D**) for this initial proof-of-concept efficacy evaluation, at the cost of a direct route-and schedule-matched comparison with nirmatrelvir.

**Figure 6.**
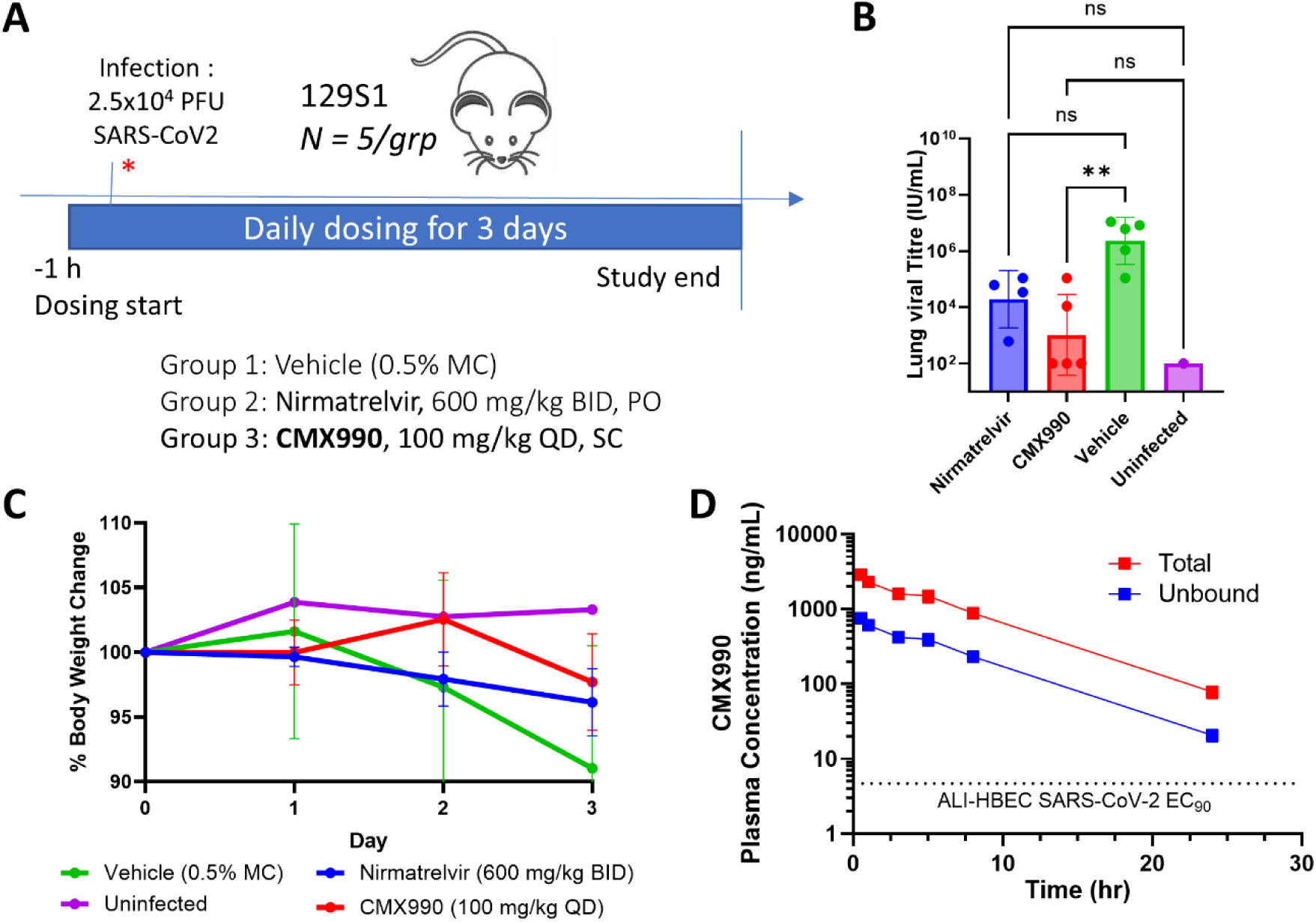
CMX990 demonstrates proof-of-concept efficacy in a murine SARS-CoV-2 infection model. (A) Dosing schematic: female 129S1 mice were intranasally infected with mouse-adapted SARS-CoV-2 and treated beginning 1 h before infection with vehicle, nirmatrelvir (600 mg/kg, PO, BID), or CMX990 (100 mg/kg, SC, QD) for 3 days. Nirmatrelvir oral dose was chosen based on prior optimized successful dose in this mouse model, and CMX990 dose was chosen based on sufficient pharmacokinetic coverage informed by mouse subcutaneous in vivo experiments. (B) Lung viral titers at study endpoint (3 dpi). CMX990 significantly reduced lung viral titers relative to vehicle (p < 0.01), while the reduction observed with nirmatrelvir relative to vehicle did not reach statistical significance (ns = non-significant). Lung viral titer data were log-transformed prior to analysis and compared by one-way ANOVA with Dunnett’s multiple-comparisons test. (C) Body weight, expressed as percent of baseline, over the 3-day dosing period; CMX990-treated animals maintained body weight comparable to the nirmatrelvir-treated group. (D) Mouse subcutaneous 100 mg/kg single-dose pharmacokinetics for CMX990 (administered as a suspension in 0.5% methylcellulose) demonstrates plasma exposure levels (total or unbound) well above the target EC_90_ threshold.

Vehicle-treated animals lost body weight over the course of infection, consistent with established disease in this model, whereas CMX990-treated mice maintained body weight equal or better than nirmatrelvir-treated group over the 3-day dosing period (**Figure 6C**). This pattern was reflected in lung viral burden at study endpoint (3 dpi, **Figure 6B**). In this study, CMX990 reduced lung viral titers relative to vehicle (p < 0.01) at a 12x lower daily dose compared to nirmatrelvir, with superior statistical significance. CMX990-treated animals showed titers closely approaching those of uninfected controls.

## Conclusion

This work provides a comprehensive characterization of the clinical stage compound CMX990, exploring its broad antiviral-breadth, structural determinants and durability of target engagement, higher barrier to resistance development, and established in vivo proof-of-concept efficacy. Across a 24-enzyme reporter panel, live-virus assays, and structural studies with SARS-CoV-1, SARS-CoV-2, and MERS-CoV M^pro^, CMX990 retains potent, mechanism-based activity across coronavirus lineages where nirmatrelvir and ensitrelvir show reduced or absent engagement outside SARS-CoV-2. This cross-lineage activity traces to a conserved covalent binding mode centered on catalytically essential residues rather than on mutable peripheral pocket contacts. The reliance on invariant catalytic architecture as well as the differentiated warhead on CMX990, offers a unifying explanation for two further findings reported here: a substantially higher barrier to resistance than nirmatrelvir or PF-00835231, without selection for the canonical E166V substitution, and durable target engagement after compound washout.^15^ Mutations resulting from CMX990 selection reflected fitness-compensatory adaptations under sustained passage pressure, rather than substitutions that directly mitigate CMX990’s covalent engagement.

These structural, resistance, and kinetic observations converge on a single mechanistic interpretation of durable covalent engagement of conserved binding site features and catalytic residues, that distinguishes CMX990 in driving unique pharmacological properties. Once-daily subcutaneous CMX990 protected against infection-associated weight loss and reduced lung viral titers to levels indistinguishable from uninfected animals in a murine SARS-CoV-2 model, providing the first *in vivo* evidence for CMX990 that these advantages translate into antiviral benefit in a live infection setting.

Collectively, these findings position CMX990 as a structurally and mechanistically validated pan-coronavirus protease inhibitor, with a profile specifically suited to pandemic preparedness: broad engagement of conserved viral machinery, resistance to rapid loss of potency under selective pressure, and durable target coverage between doses.

### Experimental Section

Unless otherwise specified, all reagents and solvents were purchased from commercial sources and used without further purification.

### Chemistry

CMX990, mCMT946, and PF-00835231 were synthesized according to procedures previously described.^5, 16^ All other compounds were purchased from commercial sources. All compounds used in this work are >95% pure by HPLC and/or NMR analyses.

### In vitro Biology

The following viral strains were obtained through BEI Resources, NIAID, NIH: Isolate HCOV-19 USA-WA1/2020 (NR-52281), Isolate hCoV-19/USA/OR-OHSU-PHL00037/2021 (Alpha Variant, NR-55461), Isolate HCOV-19/USA/PHC658/2021 (Delta Variant, NR-55691), isolate HCOV-19/USA/MD-HP20874/2021 (Omicron Variant, NR-56461), Human Coronavirus, OC43 (NR-52725), and Human Coronavirus, 229E (NR-52726). The SARS-CoV-2 assays (cell-based, enzymatic), SARS-CoV-1 assays, and MERS-CoV assays were conducted as described previously.^5, 16, 17^

### Protease Reporter Assay

Reporter-based protease inhibition data (**Figure 2**) were generated using a previously described fluorescent reporter assay^7^, with two modifications. A GFP control plasmid was used in place of the pLex expression plasmid in previous report, and P-glycoprotein inhibitor, CP-100356, was not used in current assays.

### Washout Studies

HeLa-ACE2 cells were seeded at a density of 10,000 cells per well in 96-well plates, to achieve 80-90% confluency. The following day, cells were synchronously infected with SARS-CoV-2 (WA1/2020) virus at MOI = 1 for 1 hour at 4 °C in the presence of either DMSO or compound of interest. The viral inoculum was then removed, cell monolayers washed twice with 100 µL ice-cold PBS, and compounds diluted in assay media were freshly added to each well. Compounds were tested at a single concentration corresponding to 10× their own EC_50_. Media with compound was removed either at 2, 4, or 6 h after inoculation, or retained until 10 h (no washout). For compound removal, cell monolayers were washed twice with 100 µL PBS and assay media was added to each well. At the assay end point (10 h), cell monolayers were washed with 100 µL PBS and RNA was extracted using Cells-to-CT 1-Step TaqMan Kit (Invitrogen). Viral RNA levels were quantified by RT-qPCR using the SuperScript™ III Platinum™ One-Step qRT-PCR Kit (Thermo Fisher), the 2019-nCoV N1 CDC Primers and Probe set (Integrated DNA Technologies). Human GAPD (GAPDH) was used as Endogenous Control (VIC™/MGB probe, primer limited) (Applied Biosystems) allowing for relative gene expression quantification (ddCT). Those values were normalized to DMSO control wells.

### Resistance Selection Studies

SARS-CoV-2 (WA1/2020) virus was serially passaged in HeLa-ACE2 cells in the presence of increasing concentrations of nirmatrelvir, PF-00835231, or CMX990. HeLa-ACE2 cells were seeded in 24-well plates at a density of 250,000 cells/well. Passage 1 (P1) was initiated the following day by infecting the cells with MOI = 0.01 and treating them with either DMSO or the compound of interest at a concentration of 0.5× the previously determined EC_50_ for each compound, in triplicate wells. Cultures were observed daily for evidence of CPE. Supernatants were harvested when 70-80% CPE was achieved or at the endpoint of 4 days and frozen at -80 °C for further passage. For each subsequent passage with virus, the supernatant from the previous passage was used to reinfect fresh cells; the compound concentration was increased 2-fold when active viral replication was observed at 4 days post-infection, or kept the same concentration when CPE <70% was observed. Three independent viral lineages were selected for each compound (one lineage per well), along with three vehicle (DMSO) control lineages passaged in parallel. Viruses capable of active replication at a concentration of 32× EC_50_ were obtained at P8 for nirmatrelvir, P9 for PF-00835231, and P16 for CMX990.

For phenotypic characterization, viruses were amplified for one additional passage in the absence of compound, prior to infection of assay cultures to minimize the potential for carryover of residual compound from the selection experiment. Resistant viruses were titered in HeLa-ACE2/SARS-CoV-2 cell based high content imaging assay^17^ and viral dilutions that gave 30-50% infection were used for resistance evaluation. Resistance was assessed against the compound used for selection and for cross-resistance against the other protease inhibitors in the HeLa-ACE2/SARS-CoV-2 assay, and EC_50_ fold-change was calculated between the resistant virus of interest and the EC_50_ value for the parental virus. Viral RNA was isolated from the final passage material using the QIAamp Viral RNA Mini Kit (QIAGEN), whole-genome sequencing was performed, and sequence data analyzed according to previously described methods.^18^ GenBank Accession Number: MN985325.1 was used as the reference genome.

### Protein Expression and Purification

Codon-optimized genes encoding the M^pro^ of SARS-CoV and SARS-CoV-2 for bacterial expression were purchased from Addgene (61692 and 177334, respectively). The MERS-CoV M^pro^ gene was synthesized by Integrated DNA Technologies and then cloned into the pET21b(+) vector with a C-terminal 8×His tag and a stop codon. Cells were cultured in LB medium at 37 °C to an OD₆₀₀ of ∼0.7, induced with 0.5 mM IPTG, and incubated overnight at 22 °C (SARS-CoV and MERS-CoV) and 20 °C (SARS-CoV-2). Cells were harvested by centrifugation and resuspended in lysis buffer (Buffer A: 20 mM Tris-HCl, pH 8.0, 500 mM NaCl, 5% glycerol, 0.5 mM DTT). Lysis was performed via sonication, and the clarified lysate was obtained by centrifugation at 20,000 × *g* for 45 min at 4 °C. Supernatants were loaded onto Ni-NTA affinity columns pre-equilibrated with Buffer A, washed extensively, and eluted using Buffer B (20 mM Tris-HCl, pH 8.0, 200 mM NaCl, 500 mM imidazole, 0.5 mM DTT). Eluted proteins were dialyzed against Buffer C (20 mM Tris-HCl, pH 8.0, 150 mM NaCl, 5% glycerol, 0.5 mM DTT), followed by further purification using a Superdex-200 size-exclusion column equilibrated in buffer (20 mM HEPES, pH 7.5, 150 mM NaCl, 5% glycerol, 1 mM DTT). Purified proteins were concentrated to ∼15-20 mg/mL and used immediately for crystallization.

### Crystallization, Data Collection and Structure Determination of M^pro^ – CMX990 Complexes

Freshly purified protein at 10 mg/mL in a SEC buffer was incubated with CMX990 at a 1:1 molar ratio. Crystallization was performed using the sitting-drop vapor diffusion method on our automated CrystalMation™ robotic system (Rigaku) at 20 °C at The Scripps Research Institute (TSRI), yielding diffraction-quality crystals within one week. Optimized crystallization conditions were 0.8 M ammonium sulfate, 0.1 M Bicine, pH 9.0 for SARS-CoV-1; 0.2 M magnesium chloride, 20% (w/v) polyethylene glycol 3350 for MERS-CoV; 0.2 M sodium chloride, 0.1 M Na/K phosphate, pH 6.2, and 40% PEG400 for SARS-CoV-2. Crystals were cryoprotected in their respective reservoir solution supplemented with 20% ethylene glycol as cryoprotectant and flash-cooled in liquid nitrogen. X-ray diffraction data were collected at the National Synchrotron Light Source II (NSLS-II) for SARS-CoV-1 and MERS-CoV, and at the Stanford Synchrotron Radiation Light Source (SSRL) beamline 12-2 for SARS-CoV-2. Diffraction data were indexed, integrated, and scaled using HKL-2000.^19^ Structures were solved by molecular replacement using Phaser^20^ with PDB 2A5K, 4WMD, 7Z0P as the respective MR model(s) and subjected to iterative model building and refinement using Coot^21^ and PHENIX^22^. CMX990 was modeled into the ligand-associated electron density within the M^pro^ active site (Figure S2). The final structural models were evaluated using standard crystallographic validation procedures, including MolProbity^23^. Data collection and refinement statistics are summarized in Table S1.

### In vivo Antiviral Efficacy

All animal studies were performed in an animal BSL3 facility at the Icahn School of Medicine at Mount Sinai. Procedures were conducted in accordance with institutional animal-use guidelines and protocols approved by the Mount Sinai Institutional Animal Care and Use Committee, consistent with prior SARS-CoV-2 antiviral efficacy studies.

Female 129S1 mice (7–8 week old, Jackson Laboratory Strain #002448) were intranasally infected with MA-SARS-CoV-2 at 2.5 × 10^4^ PFU per mouse in 50 µL volume. Mice were treated, beginning 1 hour prior to infection, with the test article. CMX990 was dosed subcutaneously (SC) at 100 mg/kg in 100 µL volume once daily (QD). Control groups were treated with either nirmatrelvir (MCE # PF-07321332) at 600 mg/kg in 200 µL volume PO BID, or 0.5% methylcellulose 400 cP vehicle solution PO BID. An uninfected control group was also included in the study. All formulations were created fresh each morning using 0.5% methylcellulose 400 cP solution (Sigma #M0262) vehicle. Group sizes were n = 5 for nirmatrelvir, n = 5 for CMX990, n = 5 for vehicle, and n = 1 for the uninfected control.

Body weight was measured daily following infection and expressed as percent change from baseline body weight at the time of infection. Mice were humanely euthanized using Euthasol on day 3 post infection (3 dpi). One lung was harvested for histopathology. The second lung was harvested and split in two, with one half for titers via TCID_50_ and the other half for RNA analysis.

### Pharmacokinetic (PK) Studies

Pharmacokinetic measurements for subcutaneous (SC) administration of CMX990 in CD-1 mice were made at WuXi AppTec (Shanghai, China) using the procedures previously described.^5, 16^ Three fasted animals per study group were administered subcutaneously the test article as a suspension in 0.5% methylcellulose (aqueous), at a dose concentration of 20 mg/mL. Blood samples were collected per general protocol at 0.5, 1, 3, 5, 8, and 24 h post-dosing.

The blood samples were centrifuged to obtain plasma, which was stored below −20 °C until analysis. Plasma concentrations were determined by liquid chromatography/tandem mass spectrometry (LC-MS/MS). The PK parameters were determined by non-compartmental methods using WinNonLin (v6.1 or higher version, Certara Inc., Princeton, NJ, USA).

### Lung Viral Titer Determination

Lung tissue harvested at 3 dpi was homogenized for infectious virus quantification via TCID₅₀. Vero-TMPRSS2 cells were seeded in 96-well plates at 20,000 cells/well the day before the assay. Lung samples for viral titers were homogenized in 375 µL sterile PBS and centrifuged. Supernatant was then serially diluted in infection media (DMEM with 2% FBS) 8 times and plated in triplicate onto the confluent 96-well plates. Plates were incubated at 37 °C, 5% CO_2_ for 3 days. After 3 days, plates were fixed using 10% formaldehyde, washed, and stained with 0.08% crystal violet solution for 10 minutes. Stain was then removed, plates were washed again, and cytopathic effect (CPE) on the dilution series was quantified. Titers were reported as infectious virus levels in lung homogenates.

### Statistical Analysis

Body-weight change and lung viral titers were analyzed using GraphPad Prism. Lung viral titer data were log-transformed prior to statistical analysis. Comparisons between treatment groups and vehicle were performed using ANOVA with Dunnett’s multiple-comparisons correction, matching the analysis structure used in the Prism file for the new experiment. Statistical significance was defined as P < 0.05.

## ASSOCIATED CONTENT

### Supporting Information

Additional information on protease homology and structural biology experiments.

## NotesAuthor

Authors declare no competing financial interests.

## Contributions

The manuscript was written through contributions of all authors. All authors have given approval to the final version of the manuscript.

## ACKNOWLEDGMENT

The authors would like to thank the diverse team of subject-matter experts engaged by Gates Foundation, including senior program officer Dr. Peter Warner. We also thank other Calibr-Skaggs colleagues who supported our protease inhibitor program, particularly Dr. Kelli L. Kuhen, Dr. Eugenio De Hostos, Dr. Sabine Ottilie, and Dr. Soneela Ramesh.

## ABBREVIATIONS

3CL^pro^: 3C-like protease
ACE2: angiotensin-converting enzyme 2
ALI: air-liquid interface
ANOVA: analysis of variance
BID: twice daily
BSL3: biosafety level 3
CC₅₀: half-maximal cytotoxic concentration
CCDC: Cambridge Crystallographic Data Centre
CHKV: chikungunya virus
COVID-19: coronavirus disease 2019
CoV(s): coronavirus(es)
CPE: cytopathic effect
DENV-2: dengue virus serotype 2
DMEM: Dulbecco’s Modified Eagle Medium
DMPK: drug metabolism and pharmacokinetics
DMSO: dimethyl sulfoxide
dpi: days post-infection
DPP4: dipeptidyl peptidase 4
EC₅₀: half-maximal effective concentration
EC₉₀: concentration producing 90% of maximal effect
ELA2: neutrophil elastase 2
FBS: fetal bovine serum
GFP: green fluorescent protein
HBEC: human bronchial epithelial cell(s)
HKU1: human coronavirus HKU1
hNoV: human norovirus
HRV14: human rhinovirus 14
IC₅₀: half-maximal inhibitory concentration
IF: immunofluorescence
Ki: inhibition constant
kinact: rate constant of covalent inactivation
MERS: Middle East respiratory syndrome
M^pro^: main protease (i.e., 3CL^pro^)
ND: not done
NIAID: National Institute of Allergy and Infectious Diseases
NMR: nuclear magnetic resonance
P2: amino acid position 2 of the protease recognition element
PBS: phosphate-buffered saline
PCS: Preclinical Services
PFU: plaque-forming unit(s)
PLAU: plasminogen activator urokinase
PO: per os (oral)
QD: once daily
RSV: respiratory syncytial virus
RVFV: Rift Valley fever virus
SAR: structure-activity relationship
SARS: severe acute respiratory syndrome
SC: subcutaneous(ly)
SI: selectivity index
TCID₅₀: median tissue culture infectious dose
tPA: tissue plasminogen activator
VYR: viral yield reduction
ZIKV: Zika virus

