## Supporting Information for "Pan-Coronavirus Protease Engagement and Differentiated Pharmacology of Antiviral Candidate CMX990"

### for

#### Table of Contents

**Figure S1.** Sequence alignment of SARS-CoV-1, SARS-CoV-2, and MERS-CoV main proteases ........**S2**

**Figure S2**. Electron density maps of CMX990 bound to the active site of coronavirus main proteases ...**S3**

**Table S1**.Data collection and refinement statistics for CMX990-M$^{pro}$ crystal structures………………..….**S4**

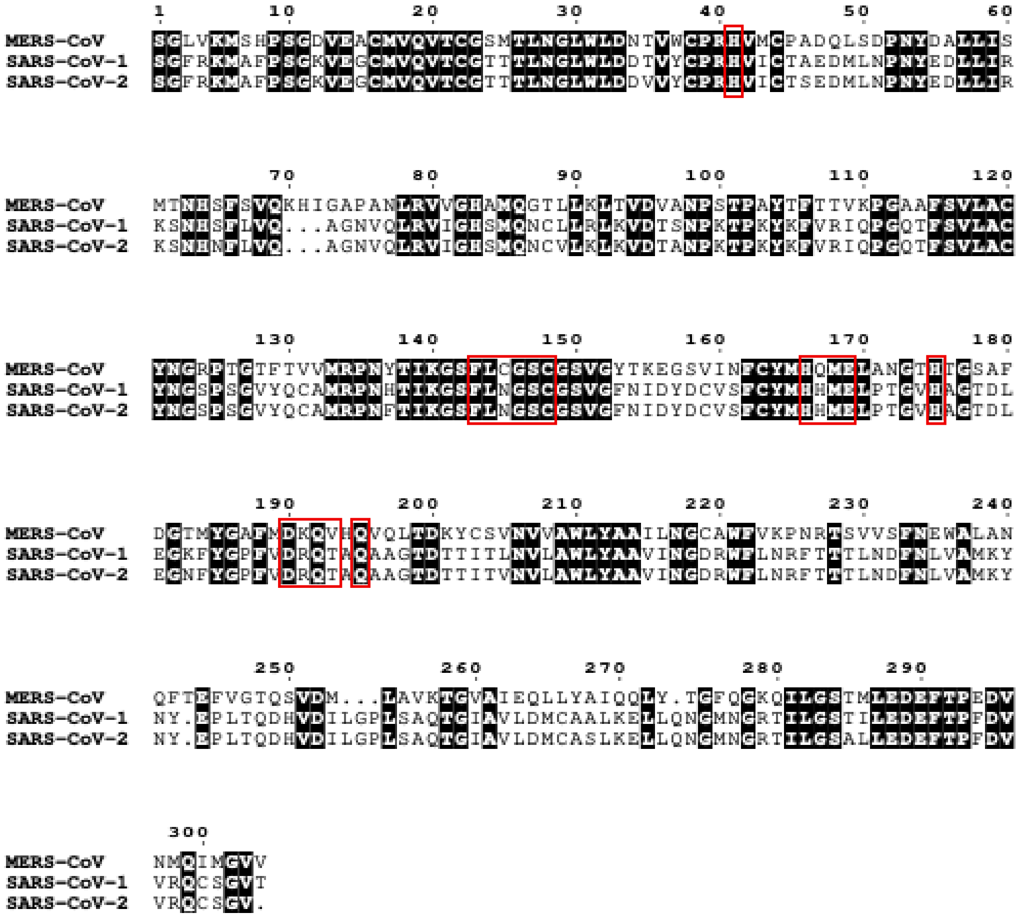

**Figure S1.** Sequence alignment of SARS-CoV-1, SARS-CoV-2, and MERS-CoV main proteases. Amino acid sequences of main protease ($M^{pro}$, nsp5) from SARS-CoV-1 (UniProt P0C6U8), SARS-CoV-2 (UniProt P0DTC1), and MERS-CoV (UniProt K9N638) were aligned to assess sequence conservation among the three structurally characterized coronavirus main proteases. Residues located within 4 Å of CMX990 in the SARS-CoV-1 $M^{pro}$–CMX990 crystal structure are highlighted with red boxes. Conserved residues are shaded according to sequence conservation. Dots indicate gaps introduced by the sequence alignment.

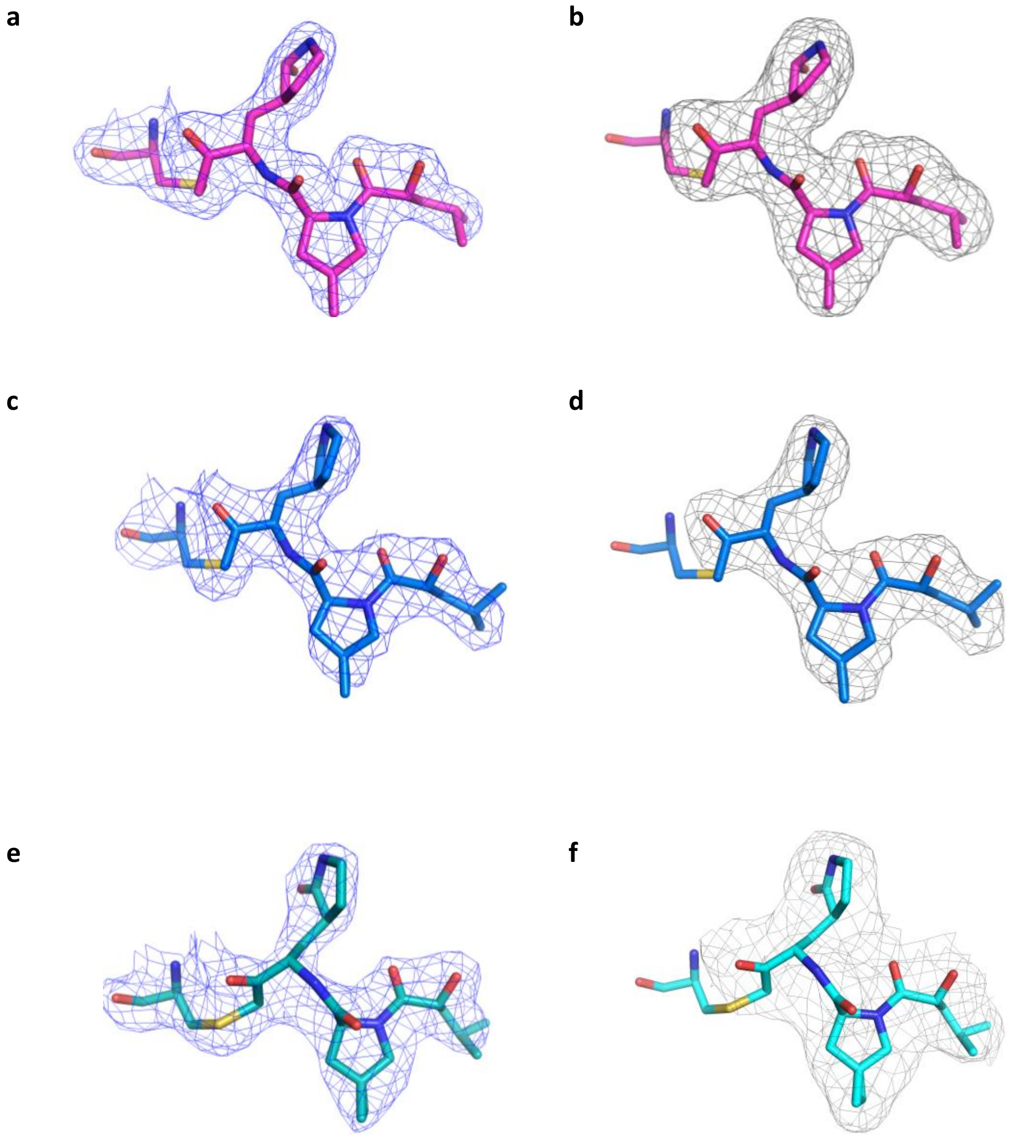

**Figure S2**. Electron density maps of CMX990 bound to the active site of coronavirus main proteases. (a, c, e) Conventional 2Fo−Fc electron density maps (blue mesh) contoured at 1.0σ for CMX990 bound to SARS-CoV-1, SARS-CoV-2, and MERS-CoV $M^{pro}$, respectively. (b, d, f) Corresponding polder omit maps (grey mesh), calculated after omission of the ligand (but not the Cys to which the ligand is attached) from the model, contoured at 2.5σ for SARS-CoV-1 and SARS-CoV-2 and 1.0σ for MERS-CoV. CMX990 is shown as sticks.

**Table S1.** Data collection and refinement statistics for CMX990-$M^{pro}$ crystal structures

| **Data Collection** | **SARS-CoV-1 +CMX990** | **MERS-CoV +CMX990** | **SARS-CoV-2 +CMX990** |
|---|---|---|---|
| Beamline | NSLS-II (17-ID-1) | NSLS-II (17-ID-1) | SSRL 12-2 |
| Wavelength (Å) | 0.92012 | 0.92012 | 0.97946 |
| Space group | $P\ 2_1\ 2_1\ 2_1$ | $P\ 1\ 2_1 1$ | $P\ 2_1\ 2_1\ 2_1$ |
| Unit cell parameters | | | |
| a, b, c (Å) | 68.0 68.3 146.9 | 62.8 112.8 92.1 | 67.7 101.0 103.4 |
| α, β, ɣ (°) | 90 90 90 | 90 91.5 90 | 90 90 90 |
| Resolution (Å)[a] | 34.1 - 2.20 (2.23 - 2.20) | 32.3 - 2.47 (2.50 - 2.47) | 50.0 - 2.54 (2.58 - 2.54) |
| Unique reflections [a] | 33,672 (1,726) | 46,145 (2,324) | 23,492 (931) |
| Redundancy [a] | 7.5 (7.2) | 7.2 (7.5) | 11.1 (6.0) |
| Completeness (%) [a] | 94.8 (100.0) | 99.7 (99.2) | 96.9 (77.8) |
| <I/σI> [a] | 7.2 (0.7) | 6.9 (1.1) | 23.3 (2.8) |
| $R_{sym}$[b] (%) [a] | 18.3 (>100) | 22.9 (>100) | 11.0 (51.0) |
| $R_{pim}$[b] (%) [a] | 7.2 (>100) | 9.1 (>100) | 3.3 (20.6) |
| $CC_{1/2}$[c] (%) [a] | 99.5 (38.5) | 99.5 (34.5) | 99.9 (97.2) |
| **Refinement Statistics** | | | |
| Resolution (Å) | 34.1 - 2.20 | 32.3 - 2.47 | 50.0 - 2.54 |
| Reflections (work) | 32,088 | 43,803 | 20,824 |
| Reflections (test) | 1,572 | 2,301 | 1,106 |
| $R_{cryst}$[d]/$R_{free}$[e] (%) | 20.8 / 25.7 | 20.7/28.6 | 20.8 / 25.5 |
| No. of copies in ASU | 2 | 4 | 2 |
| No. of atoms | 4,703 | 9,541 | 4,830 |
| $M^{pro}$ | 4,522 | 9,248 | 4,666 |
| CMX990 | 58 | 116 | 58 |
| Solvent (water) | 103 | 177 | 94 |
| DTT/GOL[f] | 20 | - | 12 (GOL) |
| Average B-values of all atoms | 58 | 48 | 37 |
| $M^{pro}$ | 59 | 47 | 37 |
| CMX990 | 55 | 48 | 36 |
| Solvent (water) | 52 | 42 | 33 |
| Wilson B-value (Å$^2$) | 47 | 37 | 37 |
| **RMSD for ideal geometry** | | | |
| Bond length (Å) | 0.008 | 0.008 | 0.002 |
| Bond Angle (°) | 0.9 | 1.0 | 0.5 |
| **Ramachandran statistics (%)**[g] | | | |
| Favored | 95 | 94 | 98 |
| Outliers | 1.2 | 0.17 | 0.7 |
| **PDB ID** | **38DQ** | **38TR** | **38TV** |

[a] Numbers in parentheses refer to the highest resolution shell

[b] $R_{sym}$ = S*hhk* S*i* | I*hkl,i* - <I*hkl*> |/ S*hkl* Si I*hkl*, *i* and $R_{pim}$ = S*hkl* (1/(n-1))1/2 S*i* | I*hkl,i* - <I*hkl*> |/S*hkl,i*, where I*hkl,i* is the scaled intensity of the ith measurement of reflection h, k, l, <I*hkl*> is the average intensity for that reflection, and *n* is the redundancy.

[c] $CC_{1/2}$ = Pearson correlation coefficient between two random half datasets.

[d] $R_{cryst}$ = S*hkl* | *Fo* – *Fc* | /S*hkl* | *Fo* | x 100, where *Fo* and *Fc* are the observed and calculated structure factors, respectively.

[e] $R_{free}$ was calculated as for $R_{cryst}$, but on a test set comprising 5% or 10% of the data excluded from refinement.

[f] DTT (dithiothreitol) and GOL (glycerol) from purification buffer

[g]From MolProbity.[1]

**References:**(1) Williams, C. J.; Headd, J. J.; Moriarty, N. W.; Prisant, M. G.; Videau, L. L.; Deis, L. N.; Verma, V.; Keedy, D. A.; Hintze, B. J.; Chen, V. B.; et al. MolProbity: More and better reference data for improved all-atom structure validation. *Protein Sci* **2018**, *27* (1), 293–315. DOI: 10.1002/pro.3330
